# Bringing the Lab to the Field: Exploring Water-Borne Corticosterone as a Physiological Indicator in Captive and Wild Common Frog Larvae (Rana temporaria)

**DOI:** 10.1101/2025.07.16.665088

**Authors:** Fabian Bartels, Katharina Ruthsatz

## Abstract

Assessing physiological condition in wild populations is important for understanding how environmental variation shapes organismal performance. Water-borne corticosterone (WB-CORT) sampling shows promise for studying amphibian stress physiology, but its ecological relevance and limitations require evaluation, particularly when methods developed under laboratory conditions are applied to wild individuals. Here, we evaluated WB-CORT sampling in common frog (*Rana temporaria*) larvae by comparing field-collected and laboratory-reared individuals. We examined whether origin influenced baseline physiological traits by modeling ontogenetic changes in WB-CORT and body mass, an integrative measure of growth. Across ontogeny, laboratory-reared larvae generally showed lower WB-CORT release and higher body mass than field-collected larvae, with distinct developmental trajectories in both traits. Next, we evaluated the sensitivity of WB-CORT and body mass to acute (48h) nitrate exposure, a widespread pollutant in amphibian breeding ponds. Nitrate treatment did not affect post-exposure WB-CORT release in either origin group. Field-collected larvae lost body mass across all treatments, including controls, whereas laboratory-reared larvae showed nitrate-specific mass loss only at 100 mg/L. Finally, we assessed whether total WB-CORT release reflected internal corticosterone by examining its relationship with tissue CORT concentrations. WB-CORT and tissue CORT were positively related in both groups, supporting WB-CORT as a minimally disruptive physiological measure under the conditions tested. Together, these findings show that WB-CORT captures biologically meaningful variation in internal corticosterone under the conditions tested, but did not detect a sustained response to the acute nitrate challenge. WB-CORT should therefore be viewed as a promising, context-dependent physiological indicator rather than a stand-alone stress biomarker.

## 1. Introduction

Understanding how organisms respond to biological, physical, or chemical environmental stressors is fundamental for predicting the impacts of global change on biodiversity. Stress can be broadly defined as a state in which environmental demands exceed an organism’s capacity to maintain or rapidly restore physiological homeostasis, thereby eliciting coordinated physiological and behavioral responses (Sapolsky et al. 2000; Romero 2004). Physiological stress responses, mediated by glucocorticoid hormones, help organisms cope with environmental challenges such as habitat degradation, climate change, pollution, and resource fluctuations (Sapolsky et al. 2000; Sapolsky 2002). These hormones, secreted by the hypothalamic–pituitary–adrenal/inter-renal (HPA/HPI) axis, regulate essential processes such as growth, energy allocation, immune function, and behavior to maintain homeostasis (Romero 2004; Crespi et al. 2013; Rollins-Smith 2017). However, when stressors are chronic, intense, or unpredictable, stress responses can lead to physiological trade-offs, including reduced growth, impaired immune function, and lower reproductive success, ultimately contributing to population declines (Gervasi & Foufopoulos 2008).

At the forefront of the global biodiversity crisis (Luedtke et al. 2023), amphibians are highly vulnerable to environmental stress owing to their permeable skin, hormone-regulated metamorphosis, and dependence on both aquatic and terrestrial habitats (rev. in Ruthsatz & Glos 2023). This exceptional sensitivity makes them widely recognized as key bioindicators of ecosystem health (Waddle 2006; Simon et al. 2011). Monitoring their physiological stress responses can therefore serve as an early warning tool (Walls & Gabor 2019), detecting signs of environmental deterioration before population declines become apparent. Despite their importance, physiological stress assessments often rely on invasive techniques like blood collection, which can induce stress, require extensive handling, and may be impractical in field settings (rev. in Ruthsatz & Rico-Millan et al. 2023). Therefore, developing, exploring, and validating non-invasive approaches for monitoring stress in wild amphibian populations is crucial for conservation efforts.

Assessing stress-related glucocorticoid activity in wild populations requires minimally invasive methods that reliably reflect endocrine state. While cortisol is the predominant glucocorticoid (GC) in primates, ungulates, and teleost fish (Schreck & Tort 2016; Suarez-Bregua et al. 2018), corticosterone (CORT) is the major GC in sauropsids (Angelier & Chastel 2009) and most amphibians (Glennemeier & Denver 2002; see Ruthsatz & Rico-Millan et al. 2023 for a review of exceptions). GC levels can be measured from a range of sample media with varying invasiveness (Madliger et al. 2018), from highly invasive (e.g., whole-body homogenates) to mildly invasive (e.g., blood, saliva, or skin swabbing), to non-invasive methods (e.g., passive urine and feces collection) (rev. in Sheriff et al. 2011). Water-borne corticosterone (WB-CORT) sampling has emerged as a promising non-invasive technique for assessing water-borne glucocorticoid release in aquatic vertebrates, including amphibians (rev. in Scott & Ellis 2007; Gabor et al. 2013; Ruthsatz et al. 2023). This method quantifies CORT released into water via gills, mucous membranes, and skin, eliminating the need for blood sampling or prolonged handling (rev. in McClelland & Woodley 2021). The water in which the specimen has been contained for a standardized time (e.g., 1–2 hours) is then collected and GCs are extracted from the water, followed by measurements through immunoassays (Burraco et al. 2015). WB-CORT allows repeated measurements across life stages while minimizing sampling stress, making it an ethically sound tool for physiological monitoring (Narayan et al. 2019; Forsburg et al. 2019). While WB-CORT has been validated in laboratory settings for some amphibian species and life-history stages (e.g., Forsburg et al. 2019; McClelland & Woodley 2021; Ruthsatz & Rico-Millan 2023), validation outcomes and approaches vary among taxa, developmental stages, and study designs, and the method has not been consistently validated across species. Its application in wild populations therefore remains limited and context dependent (but see Gabor et al. 2013; Millikin et al. 2019; Tornabene et al. 2021).

Amphibians in natural environments face multiple simultaneous stressors, including predation risk, food limitation, fluctuating temperatures, and chemical pollutants, which can influence baseline CORT levels and stress sensitivity (rev. in Ruthsatz et al. 2023; Sinai et al. 2024). At the same time, intrinsic biological factors and external environmental conditions can affect the interpretation of WB-CORT measurements (rev. in Ruthsatz et al. 2023). In amphibian larvae, CORT levels vary across ontogeny (Glennemeier & Denver 2002; Chambers et al. 2011; Ruthsatz & Rico-Millan et al. 2023), while developmental changes such as gill degeneration (Burggren & West 1982) and increasing keratinization (rev. in Schreiber & Brown 2003) may alter CORT release dynamics. External factors such as ambient temperature, skin barrier-affecting pathogens, conspecific-derived CORT, and environmental pollution can further influence WB-CORT measurements (rev. in Ruthsatz & Rico-Millan et al. 2023). Chemical contaminants are particularly relevant because they may alter physiological processes and CORT release across the skin (McClelland & Woodley 2021). Evaluating WB-CORT under such environmentally variable conditions is therefore important before extending laboratory-derived findings to field applications.

Among the various environmental stressors affecting amphibian populations, pollution is of particular concern because of its pervasive nature and potential physiological impacts (Luedtke et al. 2023; rev. in Martin et al. 2024). Many amphibians breed in water bodies associated with agricultural landscapes, where exposure to agrochemicals, including pesticides and fertilizers, is common (Brühl et al. 2013; Goessens et al. 2022). Nitrate (NO[[), a major component of fertilizers, is one of the most widespread contaminants in amphibian breeding habitats (Rouse et al. 1999; Ortiz-Santaliestra & Sparling 2007), often exceeding natural background levels in disturbed waters (<2 mg L[¹; Gomez-Isaza & Rodgers 2022). Nitrate exposure has been shown to induce a physiological stress response in amphibian larvae (Ruthsatz et al. 2022; Wang et al. 2023; McLean & Rodgers 2023), including increased CORT levels. In laboratory-reared *R. temporaria* larvae, we previously observed increased WB-CORT following nitrate exposure throughout the larval period (Ruthsatz et al. 2023). However, it remains unclear whether such WB-CORT responses observed under controlled laboratory conditions are also detectable in field-collected amphibian larvae that have developed under more heterogeneous environmental conditions. Determining whether these responses are detectable in field-collected larvae is therefore important for evaluating the transferability of laboratory-derived WB-CORT findings to field contexts.

Here, we evaluated whether WB-CORT sampling, a method previously developed and validated primarily under controlled laboratory conditions, can provide meaningful information on corticosterone variation and serve as an physiological indicator in in field-collected amphibian larvae. Using the early-breeding European common frog (*Rana temporaria*) as a model species, our study addressed two related objectives. First, we examined whether WB-CORT patterns and responses observed under laboratory conditions are transferable to field populations. To do so, we compared laboratory-reared and field-collected larvae with respect to baseline WB-CORT release and body mass across ontogeny and assessed whether the two origin groups responded similarly to short-term nitrate exposure. Second, we evaluated whether WB-CORT reflects internal corticosterone concentrations by comparing paired water-borne and tissue CORT measurements. Specifically, we aimed to: (1) examine whether field-collected larvae differ in baseline WB-CORT levels and body mass, a key integrative trait linked to growth and developmental performance, from laboratory-reared larvae; (2) assess whether the ontogenetic trajectories of WB-CORT and body mass across Gosner stages are broadly consistent between origin groups; (3) investigate whether laboratory-reared and field-collected larvae respond differently to acute nitrate exposure, potentially reflecting differences in their prior environmental histories; and (4) determine whether WB-CORT is associated with tissue CORT concentrations and therefore provides a context-dependent physiological indicator of internal endocrine state.

## 2. Material & Methods

### 2.1 Ethics statement

The experiments were conducted under permission from the Niedersächsisches Landesamt für Verbraucherschutz und Lebensmittelsicherheit, Germany (Gz. 33.19-42502-04-20/3590). Fieldwork in Lower Saxony was carried out with permits from Stadt Braunschweig (Stadt Braunschweig-Fachbereich Umwelt und Naturschutz, Richard-Wagner-Straße 1, 38 106 Braunschweig; Gz. 68.11-11.8-3.3).

### 2.2 Study species, field sampling, and experimental design

The European common frog (*R. temporaria)* is widely distributed across Europe and western Asia and occupies a broad range of natural and anthropogenic habitats, including forests, grasslands, wetlands, agricultural landscapes, and artificial water bodies (AmphibiaWeb 2026). Although currently classified as Least Concern, substantial local and regional declines have been documented in parts of Europe (Anthony & Kovács 2025), illustrating that widespread and formerly common species may undergo marked population deterioration without an immediate change in global threat status (Kyek et al. 2017; IUCN 2022). Agricultural pollution such as exposure to fertilizers is recognized as a relevant threat to the species (IUCN 2022). Its broad distribution across contrasting environments makes *R. temporaria* a useful model for examining whether physiological indicators developed under controlled conditions can detect variation relevant to field populations exposed to environmental stressors. Larval body size and developmental rate vary substantially across ontogeny and in response to environmental conditions.

To address the study hypotheses, we combined two datasets from separate breeding seasons. The 2021 dataset comprised laboratory-reared and field-collected *R. temporaria* across ontogeny and was used to examine WB-CORT release and body mass (Hypotheses 1 and 2; Fig. 1). The 2022 dataset comprised laboratory-reared and field-collected larvae sampled before and after 48-h exposure to 0, 50, or 100 mg/L nitrate and was used to assess nitrate responses and the relationship between WB-CORT and tissue CORT (Hypotheses 3 and 4; Fig. 1).Fieldwork was conducted at Kleiwiesen (52.328°N, 10.582°E), near Braunschweig, Lower Saxony, Germany, in an area surrounded by agricultural land. Pond-water nitrate concentrations were measured at animal collection in both years using an AQUA-Check 2 photometer (Söll GmbH, Germany; Table S1).

**Figure 1.**
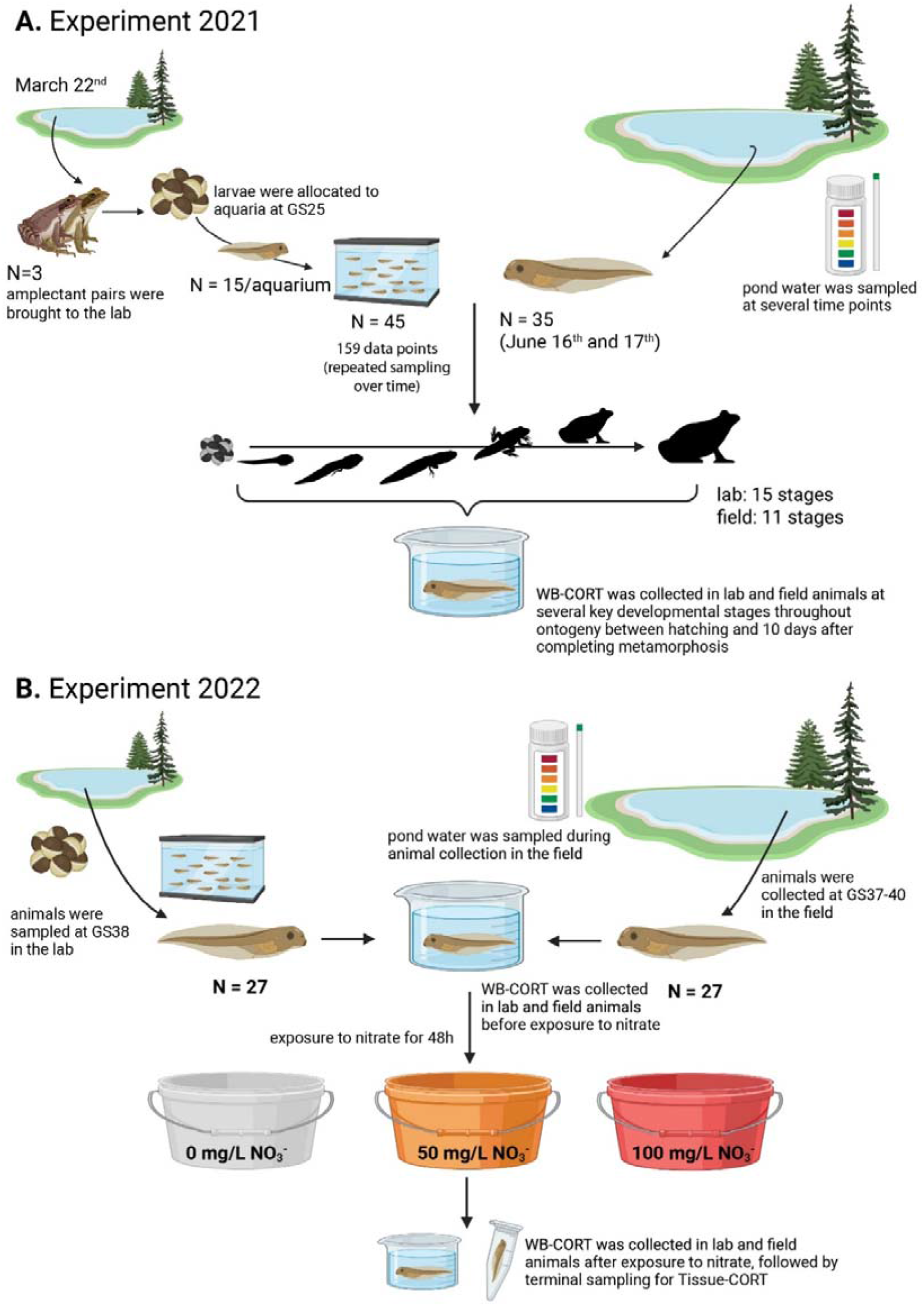
Experimental design and sampling overview. A: The 2021 experiment was used to test Hypotheses 1 and 2 by comparing ontogenetic patterns in WB-CORT release and body mass between laboratory-reared and field-collected *Rana temporaria*. B: The 2022 experiment was used to test Hypotheses 3 and 4 by assessing responses to 48 h nitrate exposure and by comparing paired post-exposure WB-CORT and tissue CORT measurements within individuals.

#### 2.2.1 2021 experiment

The 2021 dataset includes data from both laboratory and field environments (Fig. 1A). On 16 and 17 June, larvae at eleven different ontogenetic stages (Gosner stage 33 to the juvenile stage; N = 35) were collected from the field during the day for WB-CORT measurements, followed by the determination of dry-blotted body mass (see Section 2.3). Field-collected larvae hatched and developed under natural conditions in a pond at the site, and each individual was measured only once. Larvae were temporarily held in a bucket during sampling until all measurements were completed.

For lab-reared animals, the experimental design followed Ruthsatz et al. (2023) and Ruthsatz et al. (2022) but without nitrate exposure. Three amplectant pairs were collected on 22 March and transported to the Institute of Cell- and Neurobiology at TU Braunschweig, where each pair was placed in a plastic container containing 8 cm of pond water, some leaves, and branches until spawning occurred. Egg masses were collected within 12 h of oviposition, after which adults were released at the collection site. The experiment was conducted in a climate chamber (Kälte-Klimatechnik-Frauenstein GmbH, Germany) under a 14:10 h light:dark cycle at 10 ± 0.2°C during hatching and 18 ± 0.1°C from Gosner stage 25 onward (free-swimming larvae; Gosner 1960). At stage 25, larvae were transferred to three standard 12-L aquaria (15 larvae per aquarium; total N = 45), with clutches equally mixed among aquaria, resulting in an initial density of 1.66/L. Larvae were fed *ad libitum* with a 50:50 mixture of high-protein powdered fish food (Sera Micron, Sera, Germany) and *Spirulina* algae. Upon completing metamorphosis at Gosner stage 46, all juveniles (100% survival) were transferred to individual plastic containers with water to prevent desiccation and fed *ad libitum* with adult *Drosophila melanogaster* for ten days before final measurements. WB-CORT was measured at 15 individual ontogenetic stages (Gosner stage 22 to the juvenile stage), yielding 159 samples derived from 45 larvae. Larvae were repeatedly sampled throughout development; WB-CORT measured first, followed by dry-blotted body mass (see Section 2.3).

#### 2.2.2 2022 experiment

The 2022 dataset included laboratory and a field experiments to compare short-term changes in CORT and body mass following nitrate exposure in larvae that developed under either controlled laboratory conditions or natural field conditions (Fig. 1B). For the field experiment, larvae hatched and developed naturally at the Kleiwiesen pond. On 23 and 31 May 2022, 27 larvae at Gosner stages 37–40 (pro-metamorphic, with fully separated toes; Gosner 1960) were collected and temporarily held in buckets with pond water on-site until measurements were completed, without transport to the institute beforehand. For lab-reared animals, the experimental design followed Eterovick et al. (2024) and Sinai et al. (2024). Five clutches were collected at Kleiwiesen and transported to the Institute for Cell- and Neurobiology of the Technische Universität Braunschweig, where eggs were maintained at 14 ± 0.2°C in five 12-L plastic buckets containing 5 L of freshly collected pond water. At Gosner stage 25, three larvae from each clutch were transferred to each of the three 12-L aquaria containing 9 L of dechlorinated water (3 replicates × 15 larvae = 45 individuals). Larval density was 1.66 larvae/L in the beginning of the experiment. The experiment was conducted in a climate chamber under a 14:10 h light:dark cycle at 18 ± 0.1°C, and larvae were fed according to the 2021 laboratory protocol.

At Gosner stage 38 for lab-reared larvae, 27 laboratory-reared larvae were selected (nine per tank, with three from each tank assigned to each nitrate concentration), while 27 field-collected larvae were sampled at Gosner stages 37–40. WB-CORT was measured first to assess baseline CORT release, followed by determination of dry-blotted mass to the nearest 0.001 g using an electronic balance (Professional Digital Jewelry Gold Scale Balance, GandG, Germany) (see Section 2.3). Following these measurements, all 54 larvae were individually assigned to 4.4-L buckets containing 2 L of dechlorinated, ultrafiltered tap water with one of three nitrate concentrations (0, 50, or 100 mg/L). Tap- and pond-water samples (50 mL; Table S2) were collected for background WB-CORT measurements before and after nitrate exposure (see Section 2.3.1). The buckets were covered with perforated cling film for ventilation. Laboratory-reared larvae were maintained at 18 ± 0.1°C and field-collected larvae at the prevailing pond temperature (15.31 ± 2.1°C), thereby avoiding an acute temperature shift before or during WB-CORT sampling. Lab- and field-collected larvae were exposed for 48 h without water changes and were fed *ad libitum*. At the end of the exposure period, WB-CORT was measured again, followed by dry-blotted body mass (see Section 2.3). For field-collected larvae, all measurements were conducted on-site. They were then grouped into three buckets according to treatment and transported to the Institute of Cell- and Neurobiology at TU Braunschweig for tissue CORT measurements (see Section 2.3).

Nitrate treatments were prepared by adding reagent-grade sodium nitrate (NaNO[, >99% pure; Carl Roth, Germany; Oromí et al. 2009; Wang et al. 2015) to the water before introducing the larvae. Sodium nitrate is less toxic to amphibians than ammonium nitrate (NH[NO[; e.g., Johansson et al. 2001; Garriga et al. 2017). The selected nitrate concentrations fall within environmental ranges reported for German surface and groundwater (Sundermann et al. 2020) and amphibian breeding habitats (e.g., De Wijer et al. 2003; Rouse et al. 1999; Johansson et al. 2001; rev. in Sinai et al. 2024), and have previously been shown to induce increased WB-CORT in laboratory-reared larvae (Ruthsatz et al. 2023; Eterovick et al. 2024).

### 2.3 Corticosterone collection and assay

#### 2.3.1 CORT sample collection

Water-borne CORT was collected immediately after capture to limit handling stress, following the procedure of Gabor et al. (2013). Briefly, each specimen was placed in a separate 250-mL glass beaker (for larvae, metamorphs, and juveniles) containing 50 mL of ultrafiltered tap water, respectively (20 mL for juveniles to prevent drowning), for 1 hour to collect water-borne hormones. The sampling beakers were placed in a water bath (lab-reared animals; 18°C) or into the pond (field-collected animals; 15.31 ± 2.10°C for the field larvae in 2022) to maintain a constant temperature during hormone collection.

After the collection period, animals were gently removed, weighed using either a portable electronic balance in the field (Professional Digital Jewelry Gold Scale Balance, GandG, Germany) or a stationary precision balance in the lab (Sartorius A200 S, Germany). All samples were collected during the daytime, except for adult samples, which were taken between 22:00 and 02:00 h, corresponding to their peak activity period. Field-collected animals from the 2021 experiments were subsequently released at the site, whereas lab-reared animals were moved back to their aquaria and released at the Kleiwiesen site after completing metamorphosis following the requirements of the permitting authority. All water samples were stored at −20°C until processing.

In the 2022 experiment, all 54 laboratory-reared and field-collected larvae were used for paired post-exposure WB-CORT and tissue CORT measurements. Immediately after the 48-h nitrate or control exposure, WB-CORT was collected from each individual for 1 h, followed by body-mass measurement. Laboratory-reared larvae were then anesthetized in 50 mL of buffered ultrapure water containing 2 g/L tricaine methanesulfonate (MS-222; ethyl 3-aminobenzoate methanesulfonate; Sigma-Aldrich) until unresponsive to external stimuli and immediately transferred to sterile 1.5-mL tubes and snap-frozen in liquid nitrogen. For field-collected larvae, WB-CORT collection and body-mass measurements were conducted on-site, after which larvae were transported to the Institute of Cell and Neurobiology at TU Braunschweig within approximately 30 min and immediately anesthetized and snap-frozen using the same procedure. Thus, body mass was measured before anesthesia in both groups, and handling was minimized through a standardized sampling sequence conducted within a narrow time window. Handling-related stress was therefore unlikely to explain the observed variation in tissue CORT. Samples were stored at −80°C until tissue CORT extraction. These paired measurements allowed us to evaluate the relationship between post-exposure WB-CORT and tissue CORT across nitrate treatments.

To quantify background CORT, three 50-mL pond- and filtered tap-water samples were collected per water source and experiment in both years; in 2022, samples were collected before and after nitrate exposure.

#### 2.3.2 CORT sample processing

Water-borne and tissue CORT samples were processed following the procedure of Ruthsatz & Rico-Millan et al. (2023) in the respective sampling year.

Thawed water samples were first filtered through Q8 Whatman filter paper to remove suspended particles and feces, then passed through C18 solid-phase extraction columns (Oasis Vac Cartridge HLB 3 cc/60 mg, 30 μm; Waters, Inc., Switzerland) using a vacuum manifold (Visiprep Vacuum Manifold; Sigma-Aldrich, Germany). After each extraction round, the entire vacuum manifold, including the closures, was cleaned with 4 mL of HPLC-grade ethanol followed by 4 mL of nanopure water to prevent cross-contamination among samples. Columns were stored at −20°C until hormones were eluted with 4 mL of HPLC-grade methanol. Samples were transferred into 5-mL Eppendorf tubes, and methanol was evaporated using a sample concentrator (Stuart SBHCONC/1; Cole-Parmer, UK) under a fine N[stream at 45°C with a block heater (Stuart SBH130D/3; Cole-Parmer, UK). Dried samples were stored at −20°C until enzyme-immunoassay (EIA) analysis. Before analysis, samples were re-suspended in 125 μL of a 5% ethanol (95% lab grade) and 95% EIA buffer solution supplied with the DetectX Corticosterone ELISA kit (K014-H5, Arbor Assays, Ann Arbor, MI, USA) and frozen at −20°C until hormonal level measurements.

Tissue CORT samples were shipped on dry ice by overnight express to Doñana Biological Station (EBD-CSIC), Seville, Spain. Samples were randomly thawed, weighed to 0.0001 g, and homogenized in 136 × 100 mm glass tubes with 500 μL PBS buffer (AppliChem Panreac, Germany) at ∼17,000 rpm (Miccra D-1, Germany). The homogenizer was rinsed with an additional 500 μL PBS to recover residual material and cleaned with ddH[O and 96% EtOH between samples. After homogenization, 4 mL of a 30:70 petroleum ether:diethyl ether mixture (Sigma-Aldrich, Germany) was added; samples were vortexed for 60 s and centrifuged at 1800 g and 4°C for 15 min. The organic upper layer containing CORT was collected into a new 13 × 100 mm glass tube, and the extraction was repeated to maximize recovery. Pooled ether fractions were evaporated under constant nitrogen flow using a sample concentrator (Techne FSC400D; Barloworld Scientific, UK). The dried extract was resuspended in 315 μL EIA buffer (DetectX Corticosterone ELISA kit, K014-H5, Arbor Assays, Ann Arbor, MI, USA) by vortexing and incubated overnight at 4°C.

Hormone concentrations were measured using DetectX Corticosterone ELISA kits (K014-H5, Arbor Assays, USA), previously validated for *R. temporaria* (Ruthsatz et al. 2023). WB-CORT and tissue CORT were analyzed using 50-μL and 100-μL assay formats, respectively, with WB-CORT measured in duplicate and tissue CORT in triplicate on 96-well plates according to the manufacturer’s instructions. Plates were read at 450 nm using a Tecan Spark® Microplate Reader (Tecan, Switzerland) in Braunschweig and a multimode microplate reader (MB-580, Heales) in Seville. Nine plates were used in 2021 and four in 2022.

Hormone concentrations were calculated using MyAssays based on calibration standards supplied with the DetectX kit, with a new standard curve generated for each plate. Mean coefficients of variation for WB-CORT duplicates were 25.65% in 2021 and 10.92% in 2022. A pooled WB-CORT sample served as a positive control; intraplate variation was 21.64% (2021) and 15.36% (2022), and interplate variation was 29.83% and 18.34%, respectively. For tissue CORT, triplicates were retained when the coefficient of variation was ≤30.0% or the absolute mean–median difference was ≤2.5 pg; otherwise, outliers would have been removed (Ruthsatz & Rico-Millan et al. 2023). All triplicates met these criteria. Mean variation among tissue-CORT triplicates was 8.53%, with intraplate and interplate variation of 8.1% and 11.35%, respectively. Each ELISA plate included a negative control, and background corticosterone was subtracted from hormone samples. Mean R² values for the 4PLC standard curves were 0.974 (2021) and 0.992 (2022) for WB-CORT and 0.997 for tissue CORT.

WB-CORT was generally expressed as a release rate standardized by body mass and sampling duration (pg/g/h), following established practice in amphibian water-borne hormone studies. For Hypotheses 3 and 4, total WB-CORT release (pg/h) was used to avoid potential associations arising from a shared body-mass denominator. Because all WB-CORT collections lasted 1 h, the total amount released during collection corresponded to pg/h. Tissue CORT was expressed as pg/g.

### 2.4 Statistical analysis

All statistical analyses were conducted in R v.4.3.2 (R Core Team 2023). Data processing and visualization used readxl (Wickham & Bryan 2025), dplyr (Wickham et al. 2023), tidyr (Wickham & Müller 2024), ggplot2 (Wickham 2016), ggpattern (Mike FC & Davis 2022), and patchwork (Pedersen 2024); regression diagnostics and model assessment used car (Fox & Weisberg 2019) and performance (Lüdecke et al. 2021). Linear models were fitted with *lm()*. Origin-specific slopes for Hypothesis 4 were estimated with *emtrends()* from emmeans. HC3 heteroscedasticity-consistent standard errors were calculated using sandwich and used when residual heteroscedasticity persisted or in sensitivity analyses where indicated. Model assumptions were assessed visually from residual plots and statistically using Shapiro-Wilk tests (*shapiro.test()*) for normality and Breusch-Pagan tests (*bptest()*; lmtest; Zeileis & Hothorn 2002) for homoscedasticity. Model performance was evaluated using R² and diagnostic plots.

To test Hypotheses 1 and 2, mass-standardized WB-CORT release (pg/g/h) and body mass were natural-log transformed and analyzed with linear models including Origin (Field or Lab), mean-centered Gosner stage (GS_c), and their interaction. Gosner stage was treated as a continuous covariate; the juvenile stage was coded as GS 50 for analysis but is referred to as the juvenile stage in the text. Eleven WB-CORT observations from the 2021 laboratory dataset were identified as extreme outliers during diagnostics and removed (GS 25: n = 4; GS 27: n = 1; GS 35: n = 5; juvenile stage: n = 1), leaving 183 observations (148 laboratory and 35 field). For each response, the linear model was compared with models including quadratic GS terms and Origin × quadratic-GS interactions using nested model comparisons and AIC to evaluate nonlinear and origin-specific developmental trajectories. HC3 standard errors were used for inference in the selected body-mass model because residual heteroscedasticity remained and as a sensitivity analysis for WB-CORT. Because individual identity was not retained after laboratory-reared larvae were returned to their tanks, repeated observations could not be linked retrospectively and an individual-level random effect could not be fitted. Both responses were also analyzed in sensitivity analyses restricted to the developmental range shared by both origin groups (GS 33 to the juvenile stage).

To test Hypothesis 3, endocrine and body-mass responses to acute nitrate exposure were analyzed separately. Post-exposure total WB-CORT release (pg/h) was modeled as a function of pre-exposure total WB-CORT release, final body mass, Origin, nitrate treatment (0, 50, or 100 mg/L), and the Origin × Nitrate interaction. The analysis included 45 individuals with complete data, and HC3 standard errors were used because residual heteroscedasticity remained. The Origin × Nitrate interaction was evaluated with a joint robust F-test, followed by Tukey-adjusted within-origin nitrate comparisons. Body-mass change was calculated as mass before minus mass after exposure, such that positive values indicated mass loss, and analyzed in 50 individuals using a linear model with Origin, nitrate treatment, and their interaction; model assumptions were met without transformation. Nitrate treatments were compared within each origin using Tukey-adjusted estimated marginal means, and changes within each Origin × Nitrate group were tested against zero with Holm adjustment.

To address Hypothesis 4, we tested whether total WB-CORT release was associated with tissue CORT concentration and whether this relationship differed by origin. The analysis included 52 individuals with complete paired measurements; two were excluded because post-exposure WB-CORT data were unavailable. Total WB-CORT release, tissue CORT concentration (pg/g), and body mass were natural-log transformed. The model included tissue CORT, Origin, their interaction, body mass, and nitrate treatment (0, 50, or 100 mg/L), because tissue samples were collected from the same individuals immediately after the nitrate experiment. HC3 standard errors were used because residual heteroscedasticity remained after transformation, and origin-specific slopes were estimated with *emtrends()*. As a sensitivity analysis, the relationship was also tested within laboratory-reared larvae with tank identity included as a fixed blocking factor.

We used AI tools from OpenAI (ChatGPT) assist with the formulation and checking of R code for statistical analyses and data visualization. All resulting code, statistical outputs, and figures were critically reviewed and checked for accuracy by the authors. The use of AI tools was conducted in accordance with the guidelines of the German Research Foundation (Deutsche Forschungsgemeinschaft, DFG).

## 3. Results

### 3.1 Effects of origin and ontogeny on baseline WB-CORT release and body mass (Hypotheses 1 and 2)

**WB-CORT release rates -** Lab-reared larvae showed significantly lower WB-CORT levels than field-collected larvae at the mean developmental stage (β = −0.840 ± 0.238, *t* = −3.53, *p* < 0.001; Fig. 2A; Table 1). WB-CORT did not change detectably with Gosner stage in field-collected larvae (β = 0.0003 ± 0.0425, *t* = 0.008, *p* = 0.994; Fig. 2A; Table 1), whereas the significant Origin × GS interaction indicated that the ontogenetic trajectory differed between origin groups (β = −0.0998 ± 0.0437, *t* = −2.28, *p* = 0.024; Fig. 2A; Table 1). The laboratory-reared group showed a stronger decline in WB-CORT across development relative to the field-collected group. Adding quadratic Gosner-stage terms did not improve model fit, supporting the use of the linear model on the log-transformed scale. The origin difference and Origin × GS interaction remained significant when using HC3 robust standard errors and when restricting the analysis to the developmental stages shared by both groups (GS 33 – juvenile stage).

**Figure 2.**
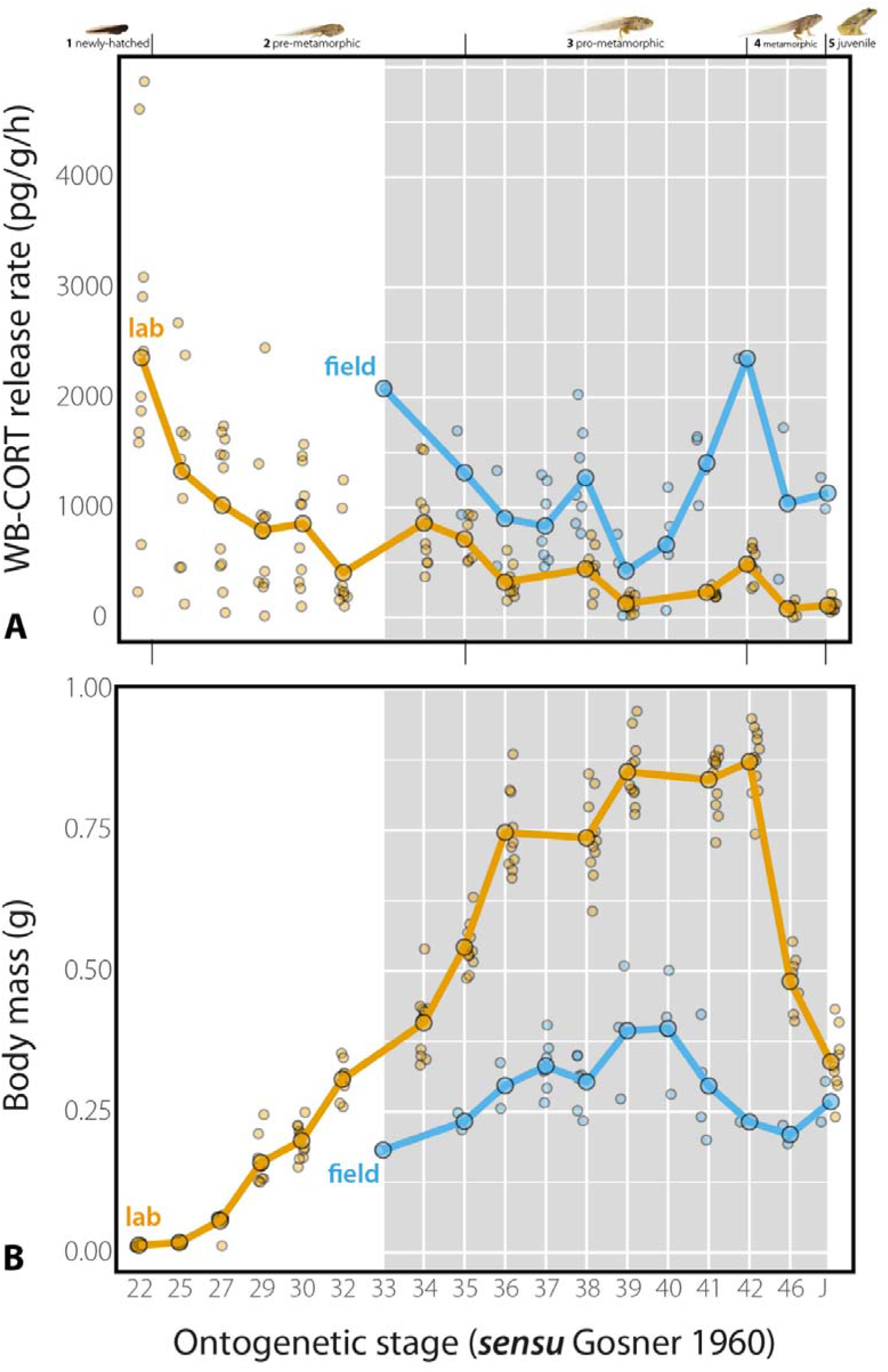
Effects of origin (laboratory-reared vs. field-collected) and ontogeny on **(A)** baseline water-borne corticosterone (WB-CORT) release rates (pg/g/h) and **(B)** body mass (g) in *Rana temporaria*. Blue symbols represent field-collected individuals and orange symbols laboratory-reared individuals. Small points show individual observations, whereas larger outlined points indicate mean values at each sampled developmental stage; lines connect these stage-specific means within each origin group. Ontogenetic stages are indicated above the figure according to Gosner (1960): **1**, newly hatched (GS 20–22); **2**, pre-metamorphic (GS 23–34); **3**, pro-metamorphic (GS 35–41); **4**, metamorphic (GS 42–46); and **5**, juvenile (>GS 46). The gray-shaded area indicates the developmental range shared by both origin groups (GS 33 to the juvenile stage). Sample sizes (laboratory/field) for WB-CORT **(A)** were: stage 1 = 11/0; stage 2 = 62/1; stage 3 = 49/29; stage 4 = 18/3; and stage 5 = 8/2. Sample sizes for body mass **(B)** were: stage 1 = 11/0; stage 2 = 67/1; stage 3 = 54/29; stage 4 = 18/3; and stage 5 = 9/2.

**Table 1.** Summary of regression model results examining the effects of larval origin (Field vs. Lab), ontogenetic stage (centered Gosner stage; GS), and acute nitrate exposure (0, 50, and 100 mg/L), as well as the relationship between tissue-based and water-borne corticosterone (CORT) in *Rana temporaria* larvae. Separate models were fitted for WB-CORT release and body mass across ontogeny (Hypotheses 1–2), post-exposure total WB-CORT release and body-mass change following 48 h of nitrate exposure (Hypothesis 3), and the relationship between tissue CORT concentration and total WB-CORT release (Hypothesis 4). Estimates represent model coefficients with associated standard errors, *t*-values, and *p*-values. Field and 0 mg/L nitrate were used as reference levels where applicable. HC3 heteroscedasticity-consistent standard errors are reported for the Hypotheses 1–2 body-mass model, the Hypothesis 3 WB-CORT model, and the Hypothesis 4 model because residual heteroscedasticity remained; conventional model-based standard errors are reported for the Hypotheses 1–2 WB-CORT model and the Hypothesis 3 body-mass model, for which model assumptions were adequately met.

| Hypothesis | Dependent variable | Fixed effects | Estimate | HC3 SE/SE | t-value | p-value |
| --- | --- | --- | --- | --- | --- | --- |
| 1 & 2 | WB-CORT release rates (pg/g/h) | Intercept (Field) | 6.725 | 0.224 | 29.918 | **<0.001** |
|  |  | Lab | -0.84 | 0.278 | -3.532 | **<0.001** |
|  |  | GS | 0.0003 | 0.043 | 0.008 | 0.994 |
|  |  | Lab × GS | -0.099 | 0.043 | -2.283 | **0.024** |
| 1 & 2 | Body mass (g) | Intercept (Field) | -1.286 | 0.087 | -14.769 | **<0.001** |
|  |  | Lab | 0.68 | 0.089 | 7.69 | **<0.001** |
|  |  | GS | 0.058 | 0.041 | 1.413 | 0.159 |
|  |  | GS<sup>2</sup> | -0.005 | 0.003 | -1.625 | 0.106 |
|  |  | Lab × GS | 0.09 | 0.041 | 2.186 | **0.03** |
|  |  | Lab × GS<sup>2</sup> | -0.007 | 0.003 | -2.43 | **0.016** |
| 3 | Post-exposure total WB-CORT release (pg/h) | Intercept (Field, 0 mg/L) | 571.289 | 305.785 | 1.868 | 0.069 |
|  |  | pre-exposure total WB-CORT (pg/h) | 0.157 | 0.205 | 0.763 | 0.45 |
|  |  | final body mass (g) | -408.795 | 744.378 | -0.549 | 0.586 |
|  |  | Lab | -167.482 | 288.462 | -0.581 | 0.565 |
|  |  | 50 mg/L | -138.386 | 117.541 | -1.177 | 0.083 |
|  |  | 100 mg/L | -68.493 | 128.35 | -0.534 | 0.597 |
|  |  | Lab × 50 mg/L | 246.303 | 138.319 | 1.781 | 0.083 |
|  |  | Lab × 100 mg/L | 305.781 | 228.419 | 1.339 | 0.189 |
| 3 | Change in body mass (g; before – after)* | Intercept (Field, 0 mg/L) | 0.076 | 0.015 | 5.193 | **<0.001** |
|  |  | Lab | -0.068 | 0.012 | -3.481 | **0.001** |
|  |  | 50 mg/L | -0.024 | 0.012 | -1.234 | 0.224 |
|  |  | 100 mg/L | -0.0315 | 0.012 | -1.607 | 0.115 |
|  |  | Lab × 50 mg/L | 0.01 | 0.028 | 0.364 | 0.718 |
|  |  | Lab × 100 mg/L | 0.117 | 0.027 | 4.370 | **<0.001** |
| 4 | ln total WB-CORT release (pg/h) | Intercept (WB-CORT) | -0.385 | 0.494 | -0.779 | 0.440 |
|  |  | ln tissue CORT | 1.119 | 0.081 | 13.841 | <0.001 |
|  |  | Lab | -1.130 | 0.784 | -1.442 | 0.156 |
|  |  | ln body mass (g) | 1.657 | 0.28 | 5.926 | <0.001 |
|  |  | 50 mg/L | -0.052 | 0.09 | -0.584 | 0.562 |
|  |  | 100 mg/L | 0.072 | 0.083 | 0.863 | 0.393 |
|  |  | ln tissue CORT ×<br>Lab | 0.206 | 0.123 | 1.68 | 0.1 |

\* Positive values indicate mass loss.

**Body mass -** Body mass showed a pronounced nonlinear ontogenetic pattern that differed between origin groups. Lab-reared larvae were significantly heavier than field-collected larvae at the mean developmental stage (β = 0.687 ± 0.089, *t* = 7.69, *p* < 0.001; Fig. 2B; Table 1). The Origin × GS (β = 0.090 ± 0.041, *t* = 2.19, *p* = 0.030; Fig. 2B; Table 1) and Origin × GS² interactions (β = −0.00761 ± 0.00313, *t* = −2.43, *p* = 0.016; Fig. 2B; Table 1) were significant using HC3 robust standard errors, indicating that the nonlinear body-mass trajectory differed between origin groups. Allowing quadratic curvature to differ between origins improved model fit (*p* = 0.0027). This difference remained evident when the analysis was restricted to the developmental stages shared between groups (GS 33– juvenile stage; Origin × GS², HC3 *p* = 0.019).

### 3.2 Origin influences mass loss, but not CORT responses, to acute nitrate exposure (Hypothesis 3)

**WB-CORT release** – Acute nitrate exposure did not produce a detectable change in total WB-CORT release (Fig. 3A; Table 1). After accounting for pre-exposure total WB-CORT release and final body mass, the Origin × Nitrate interaction was not significant (robust *F*[,[[= 1.82, *p* = 0.177; Fig. 3A; Table 1). Likewise, none of the nitrate treatments differed significantly within either field-collected or laboratory-reared larvae (all Tukey-adjusted *p* ≥ 0.349). Thus, we found no evidence that post-exposure total WB-CORT release differed among nitrate treatments in either origin group.

**Figure 3.**
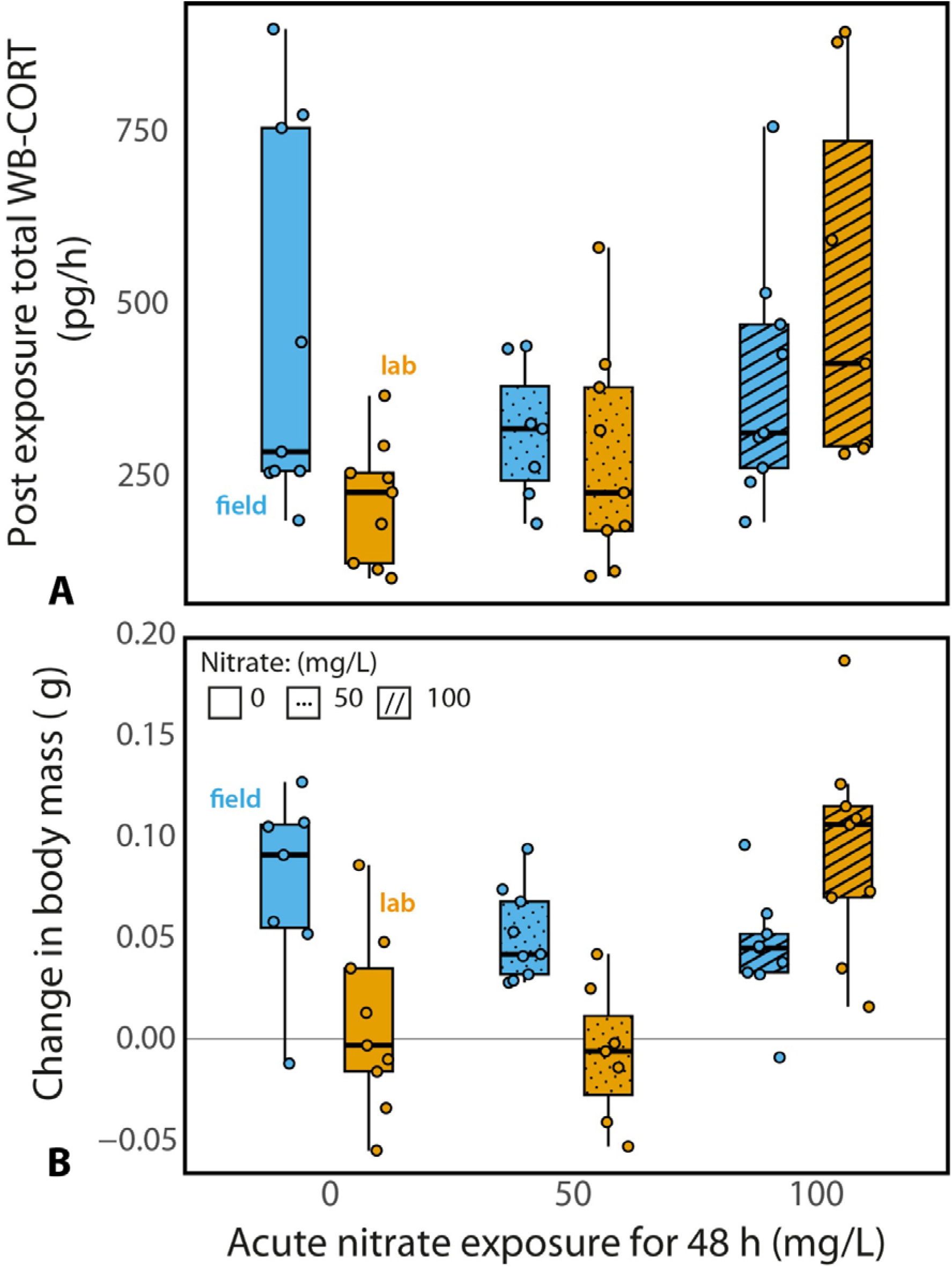
Effects of acute nitrate exposure on **(A)** post-exposure total water-borne corticosterone (WB-CORT) release (pg/h) and **(B)** change in body mass (g) in field-collected and laboratory-reared *Rana temporaria* larvae. Larvae were exposed for 48 h to 0, 50, or 100 mg/L nitrate. Blue symbols and boxes represent field-collected larvae, and orange symbols and boxes represent laboratory-reared larvae. Small points show individual observations. Boxes indicate the interquartile range, horizontal lines within boxes indicate the median, and whiskers show the range of the data. In **(B)**, body-mass change was calculated as mass before minus mass after exposure, such that positive values indicate mass loss and negative values indicate mass gain; the horizontal gray line marks zero change. For **(A)**, nitrate effects were analyzed using post-exposure total WB-CORT as the response while accounting for pre-exposure WB-CORT and final body mass. For **(B)**, body-mass change was analyzed directly.

**Body mass** – Body-mass responses differed significantly between origin groups across nitrate treatments (Origin × Nitrate: *F*[,[[= 11.99, *p* < 0.001; Fig. 3A; Table 1). Field-collected larvae lost body mass across all treatments, including the control (estimated mean loss: 0 mg/L = 0.076 g, 50 mg/L = 0.052 g, 100 mg/L = 0.045 g; all Holm-adjusted *p* ≤ 0.0037; Fig. 3A; Table 1). However, the magnitude of mass loss did not differ among nitrate treatments within field-collected larvae (all Tukey-adjusted *p* ≥ 0.253), indicating that this pattern was not nitrate-specific. In contrast, laboratory-reared larvae showed no detectable mass change at 0 or 50 mg/L, but lost an estimated 0.094 g at 100 mg/L nitrate (Holm-adjusted *p* < 0.0001). This mass loss was significantly greater than that observed at both 0 mg/L (*p* = 0.0001) and 50 mg/L (*p* < 0.0001).

### 3.3 Total WB-CORT release is positively associated with tissue CORT in both origin groups (Hypothesis 4)

Total WB-CORT release increased consistently with tissue CORT concentration in both field-collected and laboratory-reared larvae after accounting for body mass and nitrate treatment. The relationship was strong in both origin groups, with estimated slopes of 1.12 for field-collected larvae (95% CI = 0.956–1.281; Fig. 4; Table 1) and 1.32 for laboratory-reared larvae (95% CI = 1.131–1.518; both p < 0.001; Fig. 4; Table 1). Although the association appeared somewhat steeper in laboratory-reared larvae, there was no clear statistical evidence that the relationship differed between origin groups (Tissue CORT × Origin: β = 0.206 ± 0.123, t = 1.68, p = 0.100; Fig. 4; Table 1).

**Figure 4.**
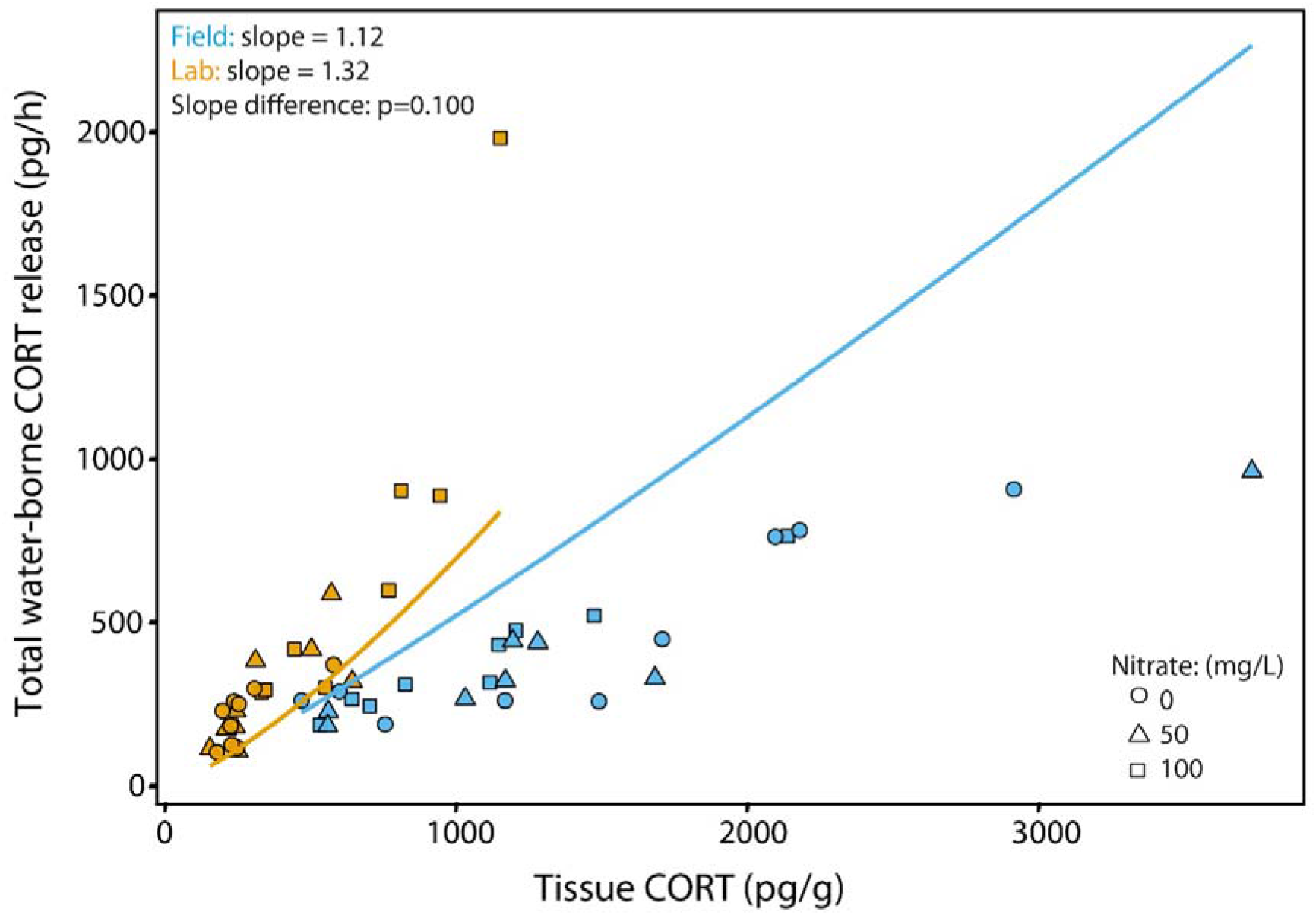
Relationship between tissue corticosterone concentration (pg/g) and total water-borne corticosterone (WB-CORT, pg/h) release in *Rana temporaria* tadpoles. Each symbol represents an individual tadpole at Gosner stages 37–40 for field-collected animals and Gosner stage 38 for laboratory-reared animals. Blue symbols indicate field-collected tadpoles and orange symbols indicate laboratory-reared tadpoles. Circles indicate 0 mg/L nitrate, triangles 50 mg/L nitrate, and squares 100 mg/L nitrate. Solid lines show model-estimated relationships between tissue CORT and total WB-CORT release for each origin group, based on the final log-transformed model and shown at median body mass and the reference nitrate treatment (0 mg/L).

## 4. Discussion

### 4.1 Origin is associated with differences in CORT levels and growth patterns in amphibian larvae (Hypotheses 1 and 2)

Understanding how baseline physiological traits vary between lab-reared and field-collected individuals is essential for evaluating the ecological relevance of stress metrics, especially when applying tools developed or validated under controlled laboratory conditions to populations exposed to natural environmental complexity (Tornabene et al. 2021). Our study revealed that origin was significantly associated with both body mass and baseline WB-CORT release in larvae of *R. temporaria*, with different ontogenetic trajectories between field-collected and lab-reared individuals. Across the developmental stages represented in both groups (GS 33 to the juvenile stage), laboratory-reared larvae generally showed lower WB-CORT release than field-collected larvae. This difference became more pronounced with advancing development because WB-CORT remained relatively stable in field-collected larvae but declined in laboratory-reared larvae. Laboratory-reared larvae were also generally heavier than field-collected larvae across the shared developmental range, although body mass followed distinct nonlinear ontogenetic trajectories in the two groups.

Unpredictable or chronic stressors can activate the HPI axis, elevating baseline glucocorticoids and diverting resources from growth to maintenance (Awkerman et al. 2024; Romero 2004; Rich & Romero 2005; Boonstra 2013). Unlike lab-reared larvae raised under stable conditions with *ad libitum* food, field-collected larvae must forage actively under environmental uncertainty, likely increasing energy expenditure and physiological demand (Moyers & Hagger 2023), which could contribute to their lower body mass (Bryant et al. 2022). However, interpretation of these origin effects should be tempered by uneven developmental stage representation in the present study, particularly among field-collected larvae, where early (GS 20–34) and late (GS 42) stages were scarce. This reflects well-known sampling challenges in natural ponds, as early stages occupy cryptic microhabitats and late-stage larvae are less detectable. Although Gosner stage was included as a covariate, the limited representation of stage extremes may have reduced our ability to detect subtle or non-linear developmental patterns, particularly for field-collected larvae. Consequently, the observed origin-specific trajectories should not be interpreted as complete ontogenetic profiles for field animals. Nevertheless, the main origin-related patterns remained evident when analyses were restricted to the developmental range shared by both groups (GS 33 to the juvenile stage). Future studies should prioritize targeted sampling across a broader and more balanced range of Gosner stages, especially during early larval development. Another potential, but untested, source of variation is exposure to ambient glucocorticoid-like compounds in the surrounding water. Environmental glucocorticoids can originate from anthropogenic sources (e.g., pharmaceuticals; Stavreva et al. 2012) or from organisms within high-density groups (Gabor et al. 2018; Tornabene et al. 2021). ELISA-based methods cannot distinguish biologically active corticosterone from cross-reactive compounds, and the resulting WB-CORT signal may therefore conflate hormone released by the focal animal with environmental glucocorticoids or chemically similar compounds. Consequently, absolute WB-CORT values obtained under field conditions should not be interpreted as direct quantitative estimates of circulating corticosterone concentrations. Rather, WB-CORT is most informative as a relative measure of endocrine activity when sampling conditions, water background, and collection procedures are standardized. Comparisons between field- and laboratory-reared animals nevertheless require particular caution because the groups were sampled in different aquatic contexts. Chronic exposure to exogenous CORT can suppress endogenous production, potentially reducing net release (Gabor et al. 2018; Tornabene et al. 2021). Amphibians absorb glucocorticoids via skin and gills (Wack et al. 2010; Middlemis Maher et al. 2013; Gabor et al. 2019), increasing susceptibility to ambient hormones. In our study, we did not quantify environmental CORT from the aquaria or verify chemical identity, so interpretation remains speculative. This highlights a methodological challenge and underscores the need for combining immunoassays with chemical analyses (e.g., LC-MS) to characterize background glucocorticoid signals in aquatic environments and their physiological relevance.

### 4.2 Acute nitrate exposure results in origin-dependent body-mass responses but no detectable WB-CORT response (Hypothesis 3)

Acute nitrate exposure did not induce a detectable change in total WB-CORT release in either field-collected or lab-reared *R. temporaria* tadpoles after accounting for pre-exposure total WB-CORT release and final body mass. Changes in body mass differed between origin groups, but the pattern observed in field-collected tadpoles was not nitrate-specific. Field-collected tadpoles lost body mass across all treatments, including controls, and the magnitude of this loss did not differ among nitrate treatments, whereas lab-reared tadpoles showed no detectable mass change at 0 or 50 mg/L but lost significant body mass at 100 mg/L nitrate. Accordingly, the mass loss observed in field-collected tadpoles cannot be attributed to nitrate exposure. It may instead reflect the combined effects of capture, handling, individual confinement, or transfer from pond water to the experimental conditions, all of which can alter stress physiology, behavior, feeding, and energy balance in amphibians (e.g., Narayan et al. 2012). However, the present design does not allow these possibilities to be distinguished. The absence of a detectable WB-CORT response after 48 h of nitrate exposure is compatible with several non-mutually exclusive explanations. First, nitrate may not have elicited a CORT response under the concentrations and exposure conditions tested. Second, a transient endocrine response may have occurred earlier during the exposure period and returned towards baseline before post-exposure sampling. Third, endocrine responsiveness may have differed among individuals or between origin groups. Animals experiencing chronically elevated glucocorticoid activity can show attenuated HPI/HPA-axis responsiveness to subsequent challenges (Romero 2004; Rich & Romero 2005; Gabor et al. 2018). Although field-collected larvae exhibited higher baseline WB-CORT levels, the present study cannot determine whether this constrained their capacity to mount an additional response.

WB-CORT represents hormone released during a standardized collection period and therefore provides a time-integrated measure rather than a direct record of rapid endocrine dynamics. This makes the method useful for repeated, minimally invasive comparisons of relative endocrine activity under standardized conditions, but brief or low-magnitude corticosterone elevations may remain undetected (Ruthsatz et al. 2022, 2023). Because WB-CORT was measured only before and after the 48-h exposure, and tissue CORT only at the end of the experiment, we could not determine whether a transient response occurred during exposure. The absence of a nitrate effect therefore indicates that no sustained difference was detectable during the post-exposure sampling period, but it does not exclude an earlier endocrine response. A further limitation is that lab-reared and field-collected larvae were maintained and sampled at their respective rearing or ambient temperatures (18°C and approximately 15°C, respectively). Although this avoided imposing an acute temperature shift, temperature itself may influence WB-CORT release and therefore cannot be excluded as a contributor to differences between the origin groups.

Together, these results show that acute nitrate exposure did not produce a detectable WB-CORT response under the conditions tested. Body-mass responses were context dependent: mass loss in field-collected tadpoles was unrelated to nitrate treatment, whereas lab-reared tadpoles showed greater mass loss only at the highest concentration. These findings emphasize the need to interpret WB-CORT alongside complementary physiological or performance-related measures, while avoiding the assumption that the absence of a WB-CORT response necessarily indicates the absence of a short-lived endocrine response.

### 4.3 WB-CORT release is associated with internal corticosterone concentrations in field-collected and laboratory-reared larvae (Hypothesis 4)

Reliable, non-invasive methods are essential for monitoring amphibian health *in situ*, especially during sensitive larval stages. WB-CORT sampling is increasingly recognized as a promising tool, with studies supporting the relationship between WB-CORT and endogenous (tissue) hormone concentrations in amphibians under laboratory conditions (e.g., Forsburg et al. 2019; McClelland & Woodley 2021; Ruthsatz & Rico-Millan et al. 2023). In the present study, we observed a strong positive relationship between total WB-CORT release and tissue CORT concentration in *R. temporaria* larvae, providing further support for the physiological relevance of this approach. Importantly, this positive relationship was evident in both field-collected and laboratory-reared larvae. It also remained strong when body mass and prior nitrate exposure were taken into account, supporting the view that WB-CORT captures biologically meaningful variation in internal corticosterone rather than merely reflecting differences in body size. Although the relationship appeared somewhat steeper in laboratory-reared larvae, we found no clear evidence that it differed between origin groups. Thus, under the conditions tested, WB-CORT appears to provide a useful, context-dependent indication of internal corticosterone variation, while not representing a direct quantitative substitute for tissue CORT.

Here, we focused on pro-metamorphic larvae, a developmental stage with naturally elevated CORT (Denver 1998; Regueira et al. 2022) and high skin permeability due to limited keratinization and functional gills (Shi 2000), facilitating transdermal release. This physiological configuration may have contributed to the strong WB–tissue CORT relationship observed in the present study. At other stages, however, this balance between internal hormone production and transdermal release may not hold, because lower CORT levels (i.e., earlier developmental stages; Denver 1998; Regueira et al. 2022), reduced skin permeability (i.e., Nishikawa et al. 1989), and the degeneration of the gills (i.e., Burggren & West 1982) could diminish WB-CORT signal strength. Field-collected larvae may also experience additional stressors (pollutants, pathogens) that alter endocrine state or hormone-release dynamics (Gabor et al. 2018; Forsburg et al. 2021). These stage- and condition-specific limitations may influence the relationship between internal CORT and water-borne hormone release and must be carefully considered when using this method in field monitoring, to avoid misinterpreting physiological baselines or stress responses (rev. in Ruthsatz & Rico-Millan et al. 2023).

### 4.4 Implications for amphibian conservation and concluding remarks

In the face of ongoing environmental change, early detection of physiological stress is critical for identifying at-risk populations before demographic declines occur (Gabor & Walls 2020). Our study highlights both the potential and the current limitations of minimally disruptive WB-CORT sampling for monitoring physiological variation in amphibian larvae. While WB-CORT allows for repeated sampling across developmental stages, its ability to detect short-term environmental responses may be limited. This was evident in our nitrate experiment, in which WB-CORT showed no detectable treatment response, whereas body mass revealed contrasting patterns between origin groups: field-collected larvae lost mass across all treatments, while laboratory-reared larvae showed nitrate-specific mass loss only at 100 mg/L. Thus, WB-CORT and body mass captured different aspects of the short-term response, highlighting the value of interpreting endocrine measures alongside complementary physiological or performance-related traits rather than in isolation. At the same time, total WB-CORT release was strongly positively associated with tissue CORT in both field-collected and laboratory-reared larvae, even after accounting for body mass and nitrate treatment. Nevertheless, because our comparison involved only one field population and one laboratory setting, the observed differences between origin groups should not be interpreted as general effects of field versus laboratory rearing. Together, these findings indicate that WB-CORT captures biologically meaningful variation in internal CORT under the conditions tested, despite showing limited sensitivity to the acute nitrate challenge. Therefore, WB-CORT represents a promising, context-dependent physiological indicator that may complement other measures in early-warning monitoring, but further validation across species, developmental stages, and environmental settings is needed before it can be applied broadly as a conservation stress biomarker.

## 5. Author contributions

**FB:** Conceptualization (lead); Methodology (equal); Data curation (equal);; Investigation (lead); Writing – original draft (equal); Writing – review and editing (equal). **KR:** Conceptualization (supporting); Methodology (equal); Data curation (equal); Formal analysis (lead); Investigation (supporting); Writing – original draft (equal); Writing – review and editing (equal). Supervision (lead); Funding raised (lead). All authors gave final approval for publication and agreed to be held accountable for the work performed therein.

## 8. Data availability

Data, including all original measurements, and codes will be deposited in Figshare under DOI:XXX after acceptance.

## 9. Statement of ethics

The authors have no ethical conflicts to disclose.

## 10 Acknowledgements

We thank Miguel Vences for his support at TU Braunschweig. We are also grateful to Paula C. Eterovick for her assistance in the field and in animal husbandry, and to Gabriele Keunecke for her help in the laboratory in Braunschweig. We sincerely appreciate Karsten Hiller for providing the facilities to conduct WB-CORT assays in the laboratory at the BRICS (Braunschweig Integrated Centre of Systems Biology) and thank Michelle-Amirah Khalil and Antonia Henne for their laboratory assistance at BRICS. Additionally, we thank Jelena Mausbach for her valuable methodological insights and advice on WB-CORT extractions and assays during protocol establishment at TU Braunschweig. Finally, we extend our gratitude to Ivan Gomez-Mestre for enabling the CORT assays for tissue CORT samples in the laboratories at the EBD-CSIC, as well as to Rafael Rico-Millan, Pablo Burraco, Monica Gutiérrez, and Francisco Miranda for their support at the EBD-CSIC.

