## Supplementary Material for "Bringing the Lab to the Field: Exploring Water-Borne Corticosterone as a Physiological Indicator in Captive and Wild Common Frog Larvae (Rana temporaria)"

Title:

**Table S1.** *Rana temporaria* natal pond and experimental water parameters (mean ± SD). Natal pond water parameters were determined at the time of animal collection. Quality of tap water used in experiments was monitored twice per week during the experiment. Measurements included nitrate (NO_3_^-^), nitrite (NO_2_^-^), ammonium (NH_4_^+^), pH, phosphate (PO_4_^3+^), copper (Cu^2+^), iron (Fe), and lead (Pb) in mg x L^-1^.

| Experimental year | Type | Water parameters | | | | | | | | |
| --- | --- | --- | --- | --- | --- | --- | --- | --- | --- | --- |
|  |  | NO_3_^-^ | NO_2_^-^ | NH_4_^+^ | PO_4_^3+^ | Cu^2+^ | Fe | Pb | pH | N |
| 2021 | Pond water | < 6 | 0.07 ± 0.01 | 0 | 0.35 ± 0.11 | <0.02 | 0.14 ± 0.02 | 0 | 6.9 ± 0.1 | 3 |
|  | Tap water used in experiments | < 6 | 0 | 0 | 0.08 ± 0.01 | <0.02 | <0.1 | 0 | 7.1 ± 0.1 | 22 |
| 2022 | Pond water | < 6 | 0.07 ± 0.02 | 0 | 0.29 ± 0.05 | <0.02 | 0.13 ± 0.01 | 0 | 6.8 ± 0.3 | 3 |
|  | Tap water used in experiments | < 6 | 0 | 0 | 0.08 ± 0.01 | <0.02 | <0.1 | 0 | 7.2 ± 0.1 | 20 |

**Table S2.** Background water-borne corticosterone (WB CORT) concentrations measured in tap and pond water samples in 2021 and in 2022 before and after nitrate exposure. Samples (50 mL) were collected as procedural blanks to verify that water sources did not contribute detectable CORT levels during the experiment. These values indicate that pond water contains high levels of CORT, suggesting that field-collected *Rana temporaria* larvae are chronically exposed to elevated exogenous CORT, in contrast to the lab-reared larvae.

| **Year** | **ID** | **Nitrate experiment** | **Origin** | **CORT (pg/ml)** | **%CV** | **CORT correct (pg)** |
| --- | --- | --- | --- | --- | --- | --- |
| 2021 | Pond 1 | NA | Field | NA | NA | NA |
| 2021 | Pond 2 | NA | Field | 150.8 | 2.32 | 18.85 |
| 2021 | Pond 3 | NA | Field | NA | NA | NA |
| 2021 | Tap water 1 | NA | Lab | 284.2 | 7.81 | 35.525 |
| 2021 | Tap water 2 | NA | Lab | NA | NA | NA |
| 2021 | Tap water 3 | NA | Lab | NA | NA | NA |
| 2022 | Pond 1 | Before | Field | 3102 | 21.1 | 387.75 |
| 2022 | Pond 2 | Before | Field | 2007 | 20.3 | 250.875 |
| 2022 | Pond 3 | Before | Field | 3224 | 2.45 | 403 |
| 2022 | Pond 4 | Before | Field | 2681 | 4.72 | 335.125 |
| 2022 | Pond 5 | Before | Field | 3554 | 12.8 | 444.25 |
| 2022 | Pond 6 | Before | Field | 3960 | 0.873 | 495 |
| 2022 | Pond 7 | After | Field | 3018 | 12.2 | 377.25 |
| 2022 | Pond 8 | After | Field | 2418 | 5.37 | 302.25 |
| 2022 | Pond 9 | After | Field | 2640 | 14.2 | 330 |
| 2022 | Tap water 1 | Before | Field | 120.5 | 19.7 | 15.0625 |
| 2022 | Tap water 2 | Before | Field | 508.7 | 0.103 | 63.5875 |
| 2022 | Tap water 3 | Before | Field | 95.31 | 37.5 | 11.91375 |
| 2022 | Tap water 4 | Before | Field | 520.1 | 43.4 | 65.0125 |
| 2022 | Tap water 5 | Before | Field | 611.9 | 0.0408 | 76.4875 |
| 2022 | Tap water 6 | Before | Field | NA | NA | NA |
| 2022 | Tap water 7 | After | Field | NA | NA | NA |
| 2022 | Tap water 8 | After | Field | 569.3 | 8.1 | 71.1625 |
| 2022 | Tap water 9 | After | Field | 653.1 | 8.13 | 81.6375 |
| 2022 | Tap water 10 | After | Field | 688.9 | 6.45 | 86.1125 |
| 2022 | Tap water 11 | After | Field | 428.7 | 11.7 | 53.5875 |
| 2022 | Tap water 12 | After | Field | 521 | 34.1 | 65.125 |
| 2022 | Tap water 1 | Before | Lab | 392.8 | 25.1 | 49.1 |
| 2022 | Tap water 2 | Before | Lab | 377 | 10.1 | 47.125 |
| 2022 | Tap water 3 | Before | Lab | NA | NA | NA |
| 2022 | Tap water 4 | After | Lab | 198.5 | 7.66 | 24.8125 |
| 2022 | Tap water 5 | After | Lab | 185.9 | 9.22 | 23.2375 |
| 2022 | Tap water 6 | After | Lab | 430.9 | 27.3 | 53.8625 |
